# Recombinant vector vaccines and within-host evolution

**DOI:** 10.1101/545087

**Authors:** James Bull, Scott L. Nuismer, Rustom Antia

## Abstract

Many recombinant vector vaccines are capable of replication within the host. They consist of a fully competent vector backbone engineered to express an antigen from a foreign transgene. From the perspective of viral replication, the transgene is not only dispensable but may even be intrinsically detrimental. Thus vaccine revertants that delete the transgene may evolve to dominate the within-host population and in doing so reduce the antigenicity of the vaccine. We apply mathematical and computational models to study this process, including the dynamics of vaccine and revertant growth plus the dynamics of innate and adaptive immunity. Although the selective basis of vaccine evolution is easy to comprehend, the immunological consequences are not. One complication is that, despite possible fitness differences between vaccine and revertant, the opportunity for vaccine evolution is limited by the short period of growth before the viral population is cleared. Even less obvious, revertant *per se* does not interfere with immunity to vaccine except as the revertant suppresses vaccine abundance; the magnitude of this interference depends on mechanisms and timing of viral suppression. Adaptive immunity targeting the foreign antigen is also a possible basis of vaccine inferiority, but it is not worsened by vaccine evolution. Overall, we find that within-host vaccine evolution can sometimes matter to the adaptive immune response targeting the foreign antigen, but even when it does matter, simple principles of vaccine design and the control of inoculum composition can largely mitigate the effects.

**Author Summary:** Recombinant vector vaccines are live replicating viruses that are engineered to carry extra genes derived from a pathogen – and these produce proteins against which we want to generate immunity. These genes may evolve to be lost during the course of replication within an individual, and there is a concern that this can severely limit the vaccine’s efficacy. The dynamics of this process are studied here with mathematical models. The potential for vaccine evolution is somewhat reduced by the short-term growth of the vaccine population before it is suppressed by the immune response. Even when within-host evolution can be a problem, the models show that increasing the vaccine inoculum size or ensuring that the inoculum is mostly pure vaccine can largely avoid the loss of immunity arising from evolution.

## 1. Introduction

Live vaccines replicate within the host. As true of any reproducing population, these within-host vaccine populations may evolve. In the absence of vaccine transmission, any within-host evolution is a dead end and might thus seem to be irrelevant to vaccine function. But if the process is fast enough, or the vaccine population replicates long enough, the vaccine population may evolve to a state where it is ineffective or virulent – either change would be bad.

The two main types of live viral vaccines are attenuated and recombinant-vector vaccines. Most live virus vaccines in current use are attenuated, their reduced virulence typically achieved by adapting the wild-type virus to a new environment (e.g. replication in a novel cell line or low temperature), with a consequent reduced replication in humans. The use of attenuated vaccines is too risky for pathogens such as HIV, and a safer alternative is to develop a live, recombinant vector vaccine where one or a few virus antigens (proteins that elicit protective immunity) are expressed from a benign virus vector.

The expected consequences of within-host evolution differ between these two types of vaccines (Table 1). Evolution of an attenuated vaccine is likely to be a reversion toward the wild-type state, the rate of this process depending heavily on vaccine design and the duration of vaccine virus replication in the host [1]. To a first approximation, reversion toward the wild-type state should lead to the vaccination more closely resembling natural infection [2], such as higher virus densities, side-effects and disease, and possibly an increased immune response. Within-host evolution of an attenuated vaccine might also predispose the virus to better transmission – also reflecting the wild-type state – but this outcome is not assured: viral adaptation to different tissues within the host may hamper growth in and dissemination from tissues important in transmission [3].

**Table 1:** Consequence of evolution for traditional live attenuated and recombinant vector vaccines

| Factor | Attenuated vaccine | Recombinant-vector vaccine |
| --- | --- | --- |
| type of evolution | reversion toward wild-type | loss of insert |
| virulence | higher virulence | little change in virulence |
| immunity | possible increase | possible reduction |
| transmission | increase | no effect or increase |

The expected consequences for evolution of a recombinant-vectored vaccine are fundamentally different [4]. In most cases, the antigen against which immunity is sought comes from a foreign transgene inserted into a competent viral vector. The vector genome carries out all viral amplification and transmission functions, and the transgene does not contribute to any process benefiting vector reproduction. From an evolutionary perspective, the transgene is both dispensable and potentially costly: selection may favor loss of the transgene and thus loss of vaccine’s ability to elicit immunity against the antigen encoded by the transgene. This evolution therefore generates something akin to infection by the wild-type vector. As vectors are typically chosen to be avirulent, vaccine evolution will result in no more than a harmless infection that does not generate immunity to the antigen encoded by the transgene.

Considerable attention has recently been given to the evolution of attenuated vaccines and designs that retard their evolution. Evolutionary stability of attenuated vaccines seems attainable by engineering designs, including the introduction of hundreds of silent codon changes, genome rearrangements, and some types of deletions [1,5,6]. Far less thought has gone into the consequences of evolution for recombinant vector vaccines or of strategies to minimize this evolution.

Although recombinant vector vaccines are not yet in widespread use, many are under development [7,8], and their success may rest on understanding within-host evolution. Here we explore how even dead-end, within-host evolution could affect the immune responses elicited by a recombinant vector vaccine and reduce its efficacy. We consider viral vaccines and focus on those that cause short-duration (acute) infections. However, the ideas we discuss apply to live vaccines of bacteria and other pathogens.

Our overall message is that while vaccine evolution may occur it is either unlikely to be a problem (i.e., compromise the generation of immunity), or it is easily mitigated. When vaccine evolution does limit the adaptive immune response, we identify ways of escaping such outcomes. Our analysis rests on mathematical models, but most results can be explained intuitively (perhaps only in hindsight), with the main results illustrated graphically; many analyses are relegated to supplementary material. Our analysis assumes that vaccines replicate within the host untill cleared by host immunity; we exclude vaccines that reproduce for just a single infection cycle (e.g., Modified Vaccinia Virus Ankara), as they have no significant opportunity for evolution.

## 2. Why the problem is not simple: preliminaries

The key question is whether evolution of the vaccine virus (henceforth just ‘vaccine’) meaningfully affects immunity to the foreign antigen encoded by the transgene (henceforth just ‘antigen’). The potential for vaccine evolution is easy to understand. Through mutation, any large vaccine population will contain mutants that inactivate or delete the foreign transgene, and those revertants will then grow amidst the vaccine. Vaccine inferiority may accrue in two different ways: the transgenic insert and its expression may intrinsically impair vaccine growth, and adaptive immunity to the foreign antigen may impair the vaccine’s growth but not the revertant’s.

It is easy to appreciate how and why the vaccine may be inferior to the revertant, and this can result in an increase in frequency of the revertant. However, the relationship between this evolution and the extent of immunity to the vaccine antigen is more complex. We thus explain some of the factors that affect how this evolution translates into a reduction in immunity to the antigen, and why in some circumstances, substantial evolution can result in little change in immunity to the antigen, while in different situations it can result in a substantial reduction.

### Duration of infection limits evolution

The revertant may have a higher growth rate because the engineered viral vaccine carries extraneous genetic cargo (termed *intrinsic* fitness differences). During viral growth, any fitness difference means that the revertant frequency will increase with time (Fig. 1). However, the vaccine gives rise to an acute infection which has a short duration, thus limiting the possible extent of evolution. The short duration is in fact the principle factor limiting evolution.

**Figure 1:**
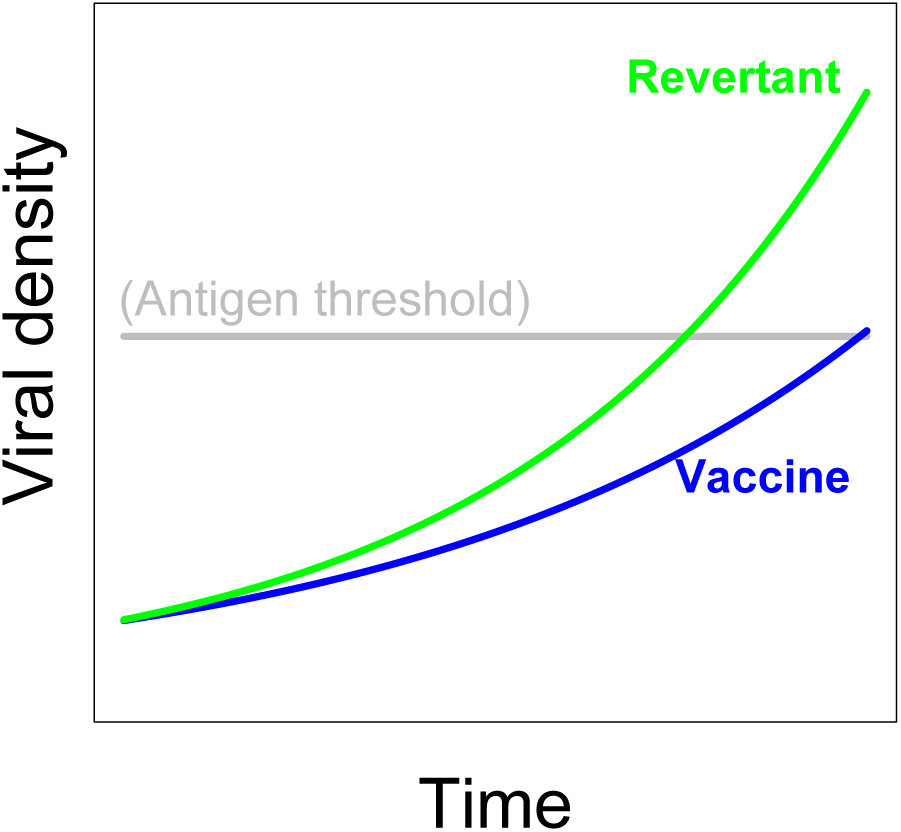
Independent growth of vaccine (blue) and revertant (green). The revertant virus has the superior growth rate, but in the absence of interference between the two, vaccine growth is unimpeded and immunity is triggered.

### Evolution versus immunity

Even more fundamentally, vaccine evolution need not reduce the immune response. If overgrowth by revertant does not interfere with vaccine growth, then antigen production is not affected (Fig. 1). Evolution affects antigen production only to the extent that revertant superiority suppresses vaccine growth and thereby suppresses antigen production.

The challenges are thus to understand (i) when and how much vaccine evolution occurs; (ii) whether and to what extent that evolution affects the abundance of vaccine virus; and (iii) the extent to which change in the vaccine abundance affects the generation of adaptive immunity against the antigen. The arguments presented above are qualitative and only superficially identify the scope of the problem. Quantitative understanding ultimately rests on analysis of mathematical models. However, as the models have many interacting processes – minimally innate immunity, adaptive immunity and intrinsic growth differences between vaccine versus revertant – we first verbally explain the biology underlying the processes that go into those models.

## 3. Bases and consequences of vaccine inferiority

An intrinsic fitness difference between the vaccine and the wild-type vector is expected (or at least not surprising) because the transgene is non-essential and has no evolutionary history with the vector genome. Thus, the insertion can be disruptive, and the resulting antigen expression may interfere with vector functions. Perhaps surprisingly, therefore, an observation of intrinsic vaccine inferiority is not necessarily the norm. Populations of recombinant viruses are commonly stable in culture, at least for a few transfers [9–20], but potentially indefinitely [21,22]. Of course, short term population retention of antigen expression may mask an underlying long term instability, so most of these observations merely set limits on the possible magnitude of inferiority. Yet even if vaccine selective ‘neutrality’ turns out to be fleeting, merely a mistaken impression from short-term observations, we will find that the phenomenon of short-term stability mirrors a solution to minimize vaccine evolution within the host.

Fig. 1 presented a hypothetical case in which evolutionary superiority of revertant did not suppress vaccine growth, hence evolution had little effect on antigen production. That process was one in which there was no interference between vaccine and revertant growth. Evolution does become important to antigen levels if vaccine and revertant interfere with either so that vaccine growth is depressed by the revertant, or if the duration of infection of the vaccine strain is reduced. In either case the revertant will then suppress antigen levels. Again, the problem is complicated by the limited duration of the infection: reduced antigen production due to vaccine evolution depends not only on interference between the two genomes but also on overall growth and the extent to which it affects the level of immunity to vaccine and vector. A mechanism that forces interference between vaccine and revertant can also limit the total amount of viral growth, thereby limiting evolution.

Evolution of vaccine versus revertant thus depends on details, in particular, the specific mechanism by which revertant interferes with vaccine growth. We describe three different mechanisms that have been proposed: innate immunity, resource limitation, and adaptive immunity to vector components. For many vaccines, each mechanism will impede revertant and vaccine equally as a collective population, thus ensuring interference.

### 3.1 Three mechanisms of vaccine-revertant interference

It was initially believed, implicitly if not explicitly, that the adaptive immune response played the dominant role in the control of viruses and other infections. In the 1990’s, Janeway and Medzhitov identified shared pathways for the control of pathogens between vertebrates and *Drosophila*, even though *Drosophila* lacks an adaptive response [23]. This led to a resurgence of interest in the role of innate immunity in the initial control of infections. Later modeling studies of influenza infections suggested yet another mechanism, that the dynamics of these infections could be largely described by simple resource limitation models, of the type used in ecology for population growth [24,25]. The realization that all three different processes might suppress viral infection led to more careful examination of the roles of different factors in the early control of acute infections [26–29]. The relative role of each mechanism in clearing infections is the basis of ongoing discussion, but it is widely accepted that the roles differ among infections by different viruses and that each mechanism is potentially important for some viruses.

### Innate immunity

There are two broad arms of immunity for suppressing vaccine growth within the host, the innate and the adaptive immune responses. Innate immunity is triggered by conserved molecules associated with pathogens [23]. Conserved structures of pathogens targeted by innate immunity include dsRNA, frequently accompanying viral replication, plus lipopolysaccharides and endotoxins of bacteria [30]. Because innate immunity involves the activation of a standing population of immune cells such as macrophages and dendritic cells, or triggering of the complement pathway, it can be elicited much more rapidly than the adaptive response; the latter requires many rounds of clonal expansion of rare antigen-specific cells to generate a population large enough to control the infection [31]. Furthermore, recent studies have shown that the innate response is required for the initial stimulation of the adaptive response [32]. Thus, innate immunity has a major role in early suppression of the viral population. Innate immunity can suppress both vector and vaccine, and it is not likely to discriminate between two genomes that differ by a single, non-essential gene (the transgene).

### Resource limitation

Another way in which virus infection can be controlled prior to the generation of adaptive immunity is resource limitation. Both the vaccine and revertant virus use the same resource (susceptible host cells). Resource limitation can control the infection if the virus depletes this resource, whereby the rate of virus output falls below its intrinsic death rate [24]. Like innate immunity, resource limitation is expected to affect vaccine and revertant similarly.

### Adaptive immunity

Adaptive immunity can be induced by the wild-type vector and the vaccine virus. Adaptive immune responses specific to antigens expressed by the wild-type vector will presumably affect the vaccine and revertant equally – because the vaccine encodes a complete vector genome, and the revertant is also a complete vector. As with the preceding pair of mechanisms, adaptive immunity common to both revertant and vaccine will operate so that revertant abundance will depress vaccine. Adaptive immunity to the vaccine antigen will be considered shortly.

All three interference mechanisms will potentially operate in any vaccinated host. With all three operating, one mechanism may take precedence over the others, simply because it is activated earlier or enforces a lower limit on viral density than the others. However, there are different stages or degrees of vaccine suppression, so an early mechanism may act to control the infection without clearing it, and another mechanism may act later to clear. Because of the delay in developing an adaptive response, viral suppression by adaptive immunity typically occurs later than effects of innate immunity or resource limitation and so might seem to be unimportant in vaccine evolution. Yet adaptive immunity may be important in clearing the vaccine following control by other mechanisms, in which case it could have an important role in vaccine evolution.

### 3.2 Adaptive immunity to the vaccine antigen may also contribute to vaccine inferiority – and feed back to inhibit itself

The preceding paragraphs omitted adaptive immunity to the antigen. By its very nature, adaptive immunity suppresses vaccine growth. But adaptive immunity to the antigen is specific to the vaccine and is thus another reason – besides intrinsic fitness effects – that the vaccine may have lower fitness than revertant. The evolutionary consequences should be the same for both types of inferiority, reducing the long term generation of antigen levels. But the interesting twist is that adaptive immunity to the antigen might feed back negatively on itself to limit its own growth – immunity against a virus is intrinsically inhibitory, so adaptive immunity against the vaccine will limit vaccine growth and thus limit antigen build-up that would fuel further immunity. One question is whether this self-inhibition is worsened with vaccine evolution.

The effect is biologically complicated because adaptive immunity to the antigen does not necessarily translate into selection against the vaccine. Selection against the vaccine *per se* operates only when adaptive immunity specifically targets the vaccine genome over the revertant genome, and this selection need not occur – either because adaptive immunity is so delayed that it is never manifest during vaccine growth, or because the antigen is physically decoupled from its genome when attacked by the adaptive response. Without imposing selection on the vaccine, antigen-directed immunity will not affect vaccine evolution.

## 4. Beyond intuition: a formal model and numerical results

We now employ quantitative models to evaluate the intuitive ideas presented above. Given the high dimensionality of the problem, we are especially interested in how well intuition works and whether generalities are observed across large regions of parameter space. A flow diagram of the elements and interactions reveals the complexity of the model (Fig. 2) and facilitates understanding the dynamical equations. *V* and *W* are the respective vaccine and revertant densities, with intrinsic growth and death rates governed by four parameters (not shown). The model also includes variables for resources (*R*), innate immunity (*Z*), adaptive immunity to vector (*Y*), and adaptive immunity to antigen (*X*) that are both influenced by and influence *V* and *W* (Fig 2). In the following sections, we explore the dynamics of these interactions with simulations and present results graphically. Equations and parameter values are provided in the Appendix. Resource limitation and innate immunity yield qualitatively similar results, so trials with resource limitation are not illustrated in the main text.

**Figure 2:**
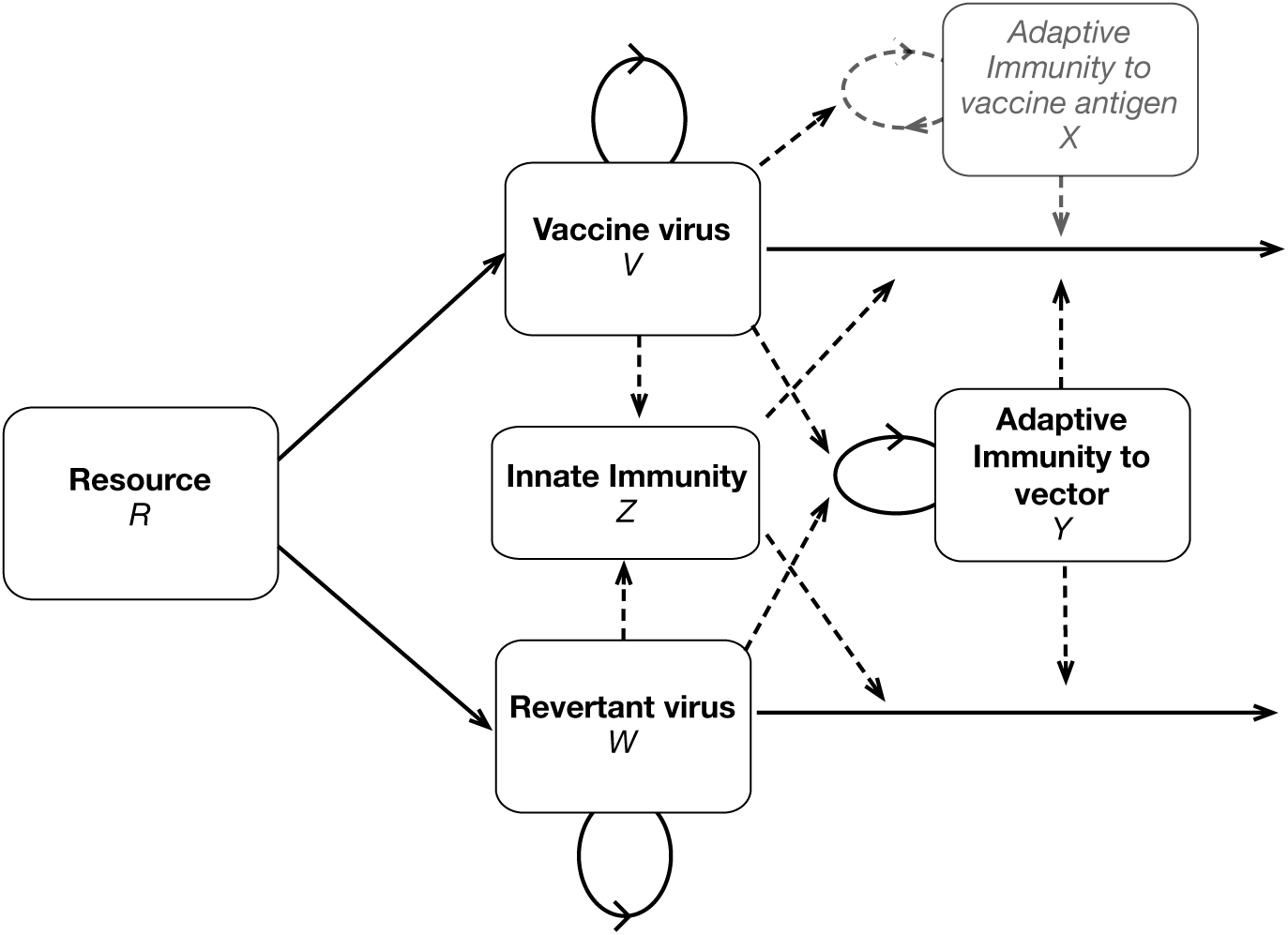
Diagram of model processes and interactions. This figure gives all the processes in the full model that includes resource limitation with innate and adaptive immunity. Solid lines represent variables (V, W, R, Z, X, and Y) and dashed lines represent influences. Note that only the top-most box in gray, the specific immune response to the vaccine antigen, acts differentially on the vaccine vs revertant virus. Not all of these components are included in each iteration of the model.

The models assist us by forcing us to specify assumptions for how the viruses and immunity interact, and by allowing us to rigorously explore outcomes in different scenarios. However, there is uncertainty in the model structure, many parameter values are unknown, and different viruses will behave somewhat differently. Consequently, we focus on broad generalities that arise from many simulations and illustrate these for a few specific cases, reserving the supplement for further details. The presentation below briefly discusses the individual dynamics of individual trials for illustration but then moves to plots that reveal differences in outcomes as the key parameters are changed. The model used here incorporates the structure of earlier models used to describe immune responses [33–35]; parameter values used here were chosen as described in some of these earlier studies.

### 4.1 Evolution from intrinsic fitness effects can matter

In the trials used for illustration, we allow innate immunity to control the infection and adaptive immunity to cause final clearance. Such a scenario might correspond to the dynamics of *Listeria* infection of mice [31], or the early dynamics of SIV infections [36]. To get a sense of the full dynamics in the model, we show the time course of dynamics for the different variables (Fig. 3). The left panel plots the dynamics of virus and immunity in the absence of evolution (revertant absent). The right panel plots two extremes of vaccine evolution, one slight (solid curves), one strong (dotted curves); vaccine evolution is enhanced by increasing the mutation rate, the fitness of revertant (*c*) and the initial revertant abundance. The effect of evolution is seen from a comparison of the dashed and solid curves on the right with each other and a comparison of those curves with the left panel.

**Figure 3:**
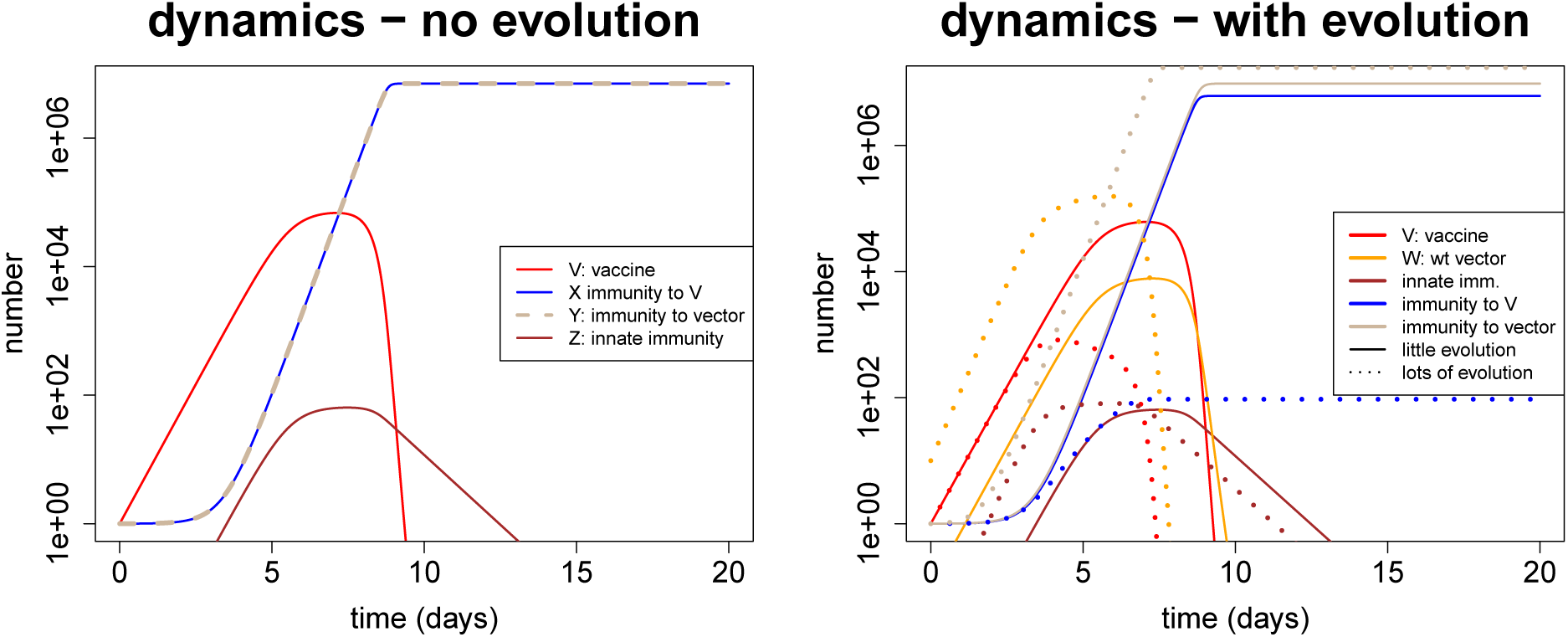
Representative dynamics contrasting vaccine evolution with no evolution. The trials are parameterized so that virus is controlled by innate immunity with final clearance due to adaptive immunity. (Left) The dynamics of virus and immunity are shown in the absence of revertant (i.e. no evolution). (Right) The revertant is included, but at two different levels. The solid lines correspond to little evolution: the vaccine has a small cost (intrinsic cost =1%, initial level of W is 0.1 that of initial vaccine, and the mutation rate is 10^−^6 per day). The dotted lines correspond to major evolution: the vaccine has a 20% intrinsic cost, the mutation rate is 10^−3^, and the initial level of the revertant is 10 fold that of the vaccine.

Comparing the cases of no evolution with little evolution, the revertant virus does not significantly affect the dynamics of the vaccine virus or immunity to the vaccine virus (red solid lines in left and right panels are similar). However, with parameters that result in considerable evolution (high mutation rate, high initial frequency and large growth advantage for the revertant virus), the vaccine virus is suppressed and cleared earlier, reducing cumulative lifetime production of vaccine virus, and thus of vaccine antigen and of immunity to the vaccine antigen.

Illustrations of dynamics from individual trials convey many details. However, without a specific empirical basis for the parameter values chosen, the details have little assured relevance. We therefore provide contour plots that allow easy comparison of many different trials (Fig. 4). These graphs show the cumulative vaccine load (left panel) and final level of immunity to vaccine (right) as a function of initial revertant frequencies and selective advantage of the revertant (*c*). A strong correspondence exists between virus load and the level of immunity generated, as is observed following infection [37]. (Subsequent figures therefore illustrate the level of immunity.) The initial composition of the inoculum matters somewhat more to the adaptive response than does the intrinsic cost of the vaccine (as evident by the contours being closer to vertical rather than horizontal). When the inoculum is mostly vaccine and revertant fitness is not high, evolution has little effect on viral load or final level of immunity (i.e., the lower left of each panel has a broad area of one color) – because of the short duration of infection. Over longer periods of time, the selective advantage of the revertant plays an increasing role in evolution.

**Figure 4:**
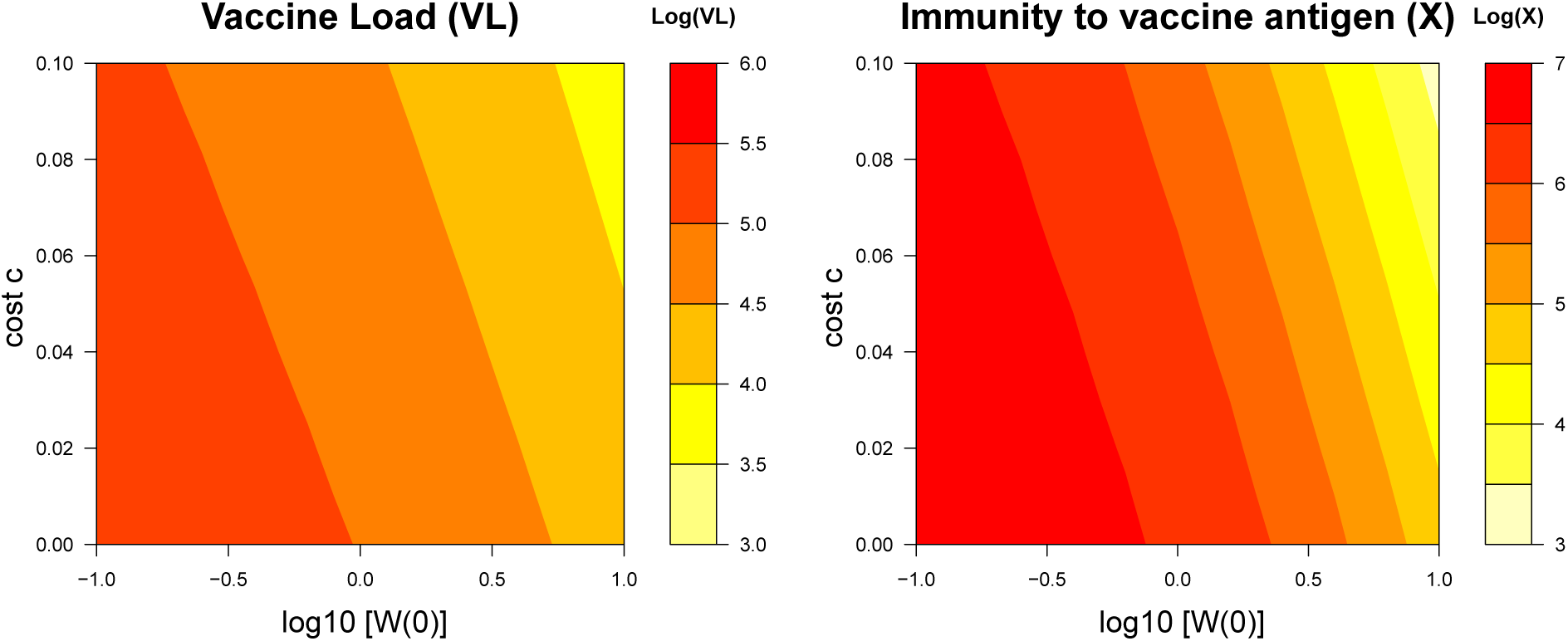
Viral load and the level of immunity to the vaccine antigen depend on evolution. The extent of evolutionary change depends largely on two parameters, the initial abundance of the revertant virus (plotted on the x-axis) and the growth advantage of the revertant (plotted on the y-axis). The heat maps show how, as the extent of evolutionary change increases (as we move to the right or up), there is a reduction in the viral load of the vaccine (defined as ∫ *V dt*, left panel) and in the magnitude of immunity to the vaccine antigen (*X*, right panel). The initial amount of vaccine virus is always V(0)=1 (log =0). Note that the graphs span high levels of revertant in the inoculum that should be easily avoided (log W(0) > 1) – if the researcher is alert to the possibility. Parameters as in SI table with: no evolution scenario having W(0)=*µ*=0 (left panel); low evolution scenario having W(0)= 0.01 V(0), *c* = 0.01, *µ* = 10^−6^; and high evolution having W(0)= 10 V(0), *c* = 0.2, *µ* = 10^−3^.

### 4.2 Vaccine evolution driven by adaptive immunity

We focus on infections of short duration. Factors that limit the duration of infection include resource limitation, and innate and adaptive immunity. For the most part these factors act equally against vaccine and revertant virus. Only one factor, adaptive immunity to the vaccine antigen (*X*), acts specifically on the vaccine virus and not the revertant. Intuition suggests that this adaptive immunity to the antigen can potentially suppress the vaccine’s growth and give an advantage to the revertant. As with intrinsic fitness costs, this selection might feed back to limit vaccine growth and thus limit the development of further immunity by allowing revertant to grow and interfere with vaccine. This section considers whether these arguments are supported by the model.

Any real vaccine that elicits immunity against the antigen may also experience an intrinsic fitness cost. The effect of immunity on evolution would then be confounded with the effect of intrinsic fitness effects on evolution, making it difficult to isolate one from the other. The models do not face this problem, however. They can be parameterized so that the only possible selection against the vaccine comes from immunity (by setting *c* = 0). Vaccine populations can also be freed of revertant by omitting revertant from the inoculum and setting the mutation rate to 0. Thus, we can measure the effect of adaptive immunity on vaccine growth from trials that lack revertant and then compare those results with trials that include revertant.

There are several background points to note about the model structure. First, adaptive immunity specific to vaccine (*X*) develops at a rate proportional to the vaccine abundance (*V*) and parameters *s* and *φ_X_*. In contrast the impairment of vaccine growth depends on the level of immunity (*X*) and the parameter (*k_X_*). Thus, immunity can develop even when there is little or no impairment, i.e., when *k_X_ →* 0. Second, adaptive immunity to the vector (*Y*) develops according to its own parameter (*φ_Y_*) in response to vaccine plus revertant abundance (*X* + *Y*), and it impairs both vaccine and revertant growth equally by parameter *k_Y_*. When revertant is present, it will increase the level of immunity to vector backbone/revertant but not directly affect immunity specific to the vaccine. This immunity will result in faster clearance of both revertant and vaccine, and this results in decreased immunity to the antigen.

Trials were run that contrasted revertant absence versus revertant introduced at 75% of the inoculum – no evolution versus evolution, respectively (Fig. 5). Absence of the revertant is the baseline against which the effect of evolution can be compared. The horizontal axis varies *k_X_*, the parameter for impairment specific to vaccine, and the vertical axis varies *k_Y_*, impairment to vector, which affects vaccine and revertant equally. In both panels, increasing impairment against vaccine leads to lower levels of immunity to the vaccine – this is the self-limiting effect of adaptive immunity, which exists even in the absence of evolution. As expected, impairment of vaccine by immunity to vector is also found.

**Figure 5:**
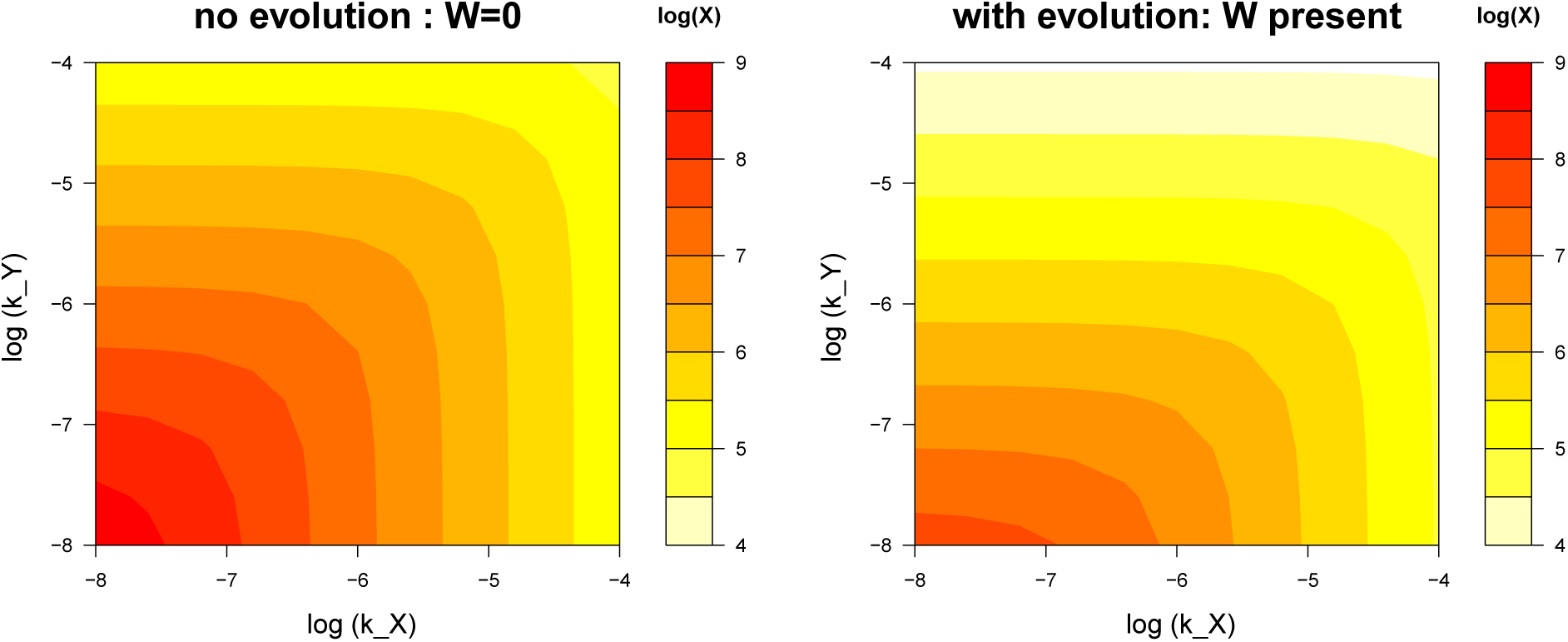
Effect of evolution on the suppression of immunity by impairment parameters. The final level of immunity to the vaccine antigen depends heavily on the parameters *k_X_* and *k_Y_* – which describe how immunity to the vaccine and revertant affect virus replication. The left plot considers the absence of revertant, hence no evolution. The right panel introduces revertant at 3/4 the inoculum, with the same total inoculum size as in the left panel. The revertant reduces immunity (*X*), but the effect of increasing *k_X_* is not made worse by the revertant. Intrinsic fitness differences are absent; mutation of vaccine to revertant is set to 0.

A large effect of evolution on vaccine immunity is evident by comparing the left panel (no evolution) and right panel (evolution): introduction of revertant reduces the level of immunity against vaccine on the order of 10-fold. For the evolution panel, the revertant is 3/4 the inoculum and has no intrinsic advantage over vaccine; inoculum size is unchanged. All loss of immunity against vaccine is thus due to revertant in the inoculum and any selective effect from immunity against vaccine.

A question motivating this analysis was one step deeper in the complexity of these effects: does the self-limiting effect of adaptive immunity worsen with evolution? This question can be answered by comparing the self-inhibitory effect between left and right panels as *k_X_* is increased. It is seen that the self-inhibitory effect is actually somewhat reduced by the revertant. The revertant lowers the response overall, but when correcting for that difference, the effect of increasing *k_X_* is weaker in the right panel than in the left. We attribute this weakening of self-limitation as due to the same effect in Fig. 1: revertant presence becomes irrelevant as more of the adaptive response to vaccine is controlled by immunity to the vaccine antigen rather than vector – revertant is interfering less.

In sum, therefore, immunity to the vaccine (*X*) is reduced by itself and by evolution (presence of revertant). The self-limiting effect of anti-vaccine immunity depends heavily on the impairment parameter. The two effects do not interact to make the problem worse than from their separate effects.

### 4.3 Optimizing the efficacy of a recombinant vector vaccine to avoid effects of evolution

Vaccine design can affect the level of immunity specific to its recombinant antigen. This section briefly consider factors that affect the efficacy of a recombinant vector vaccine in the absence of evolution, then turns to vaccine designs and administration that improve vaccine efficacy in the presence of evolution.

An ideal recombinant vector vaccine would have the following properties. First it should elicit an immune response that rapidly clears the pathogen (i.e. the rate constant for clearance of the pathogen, call it *k_P_*, is high). Second, the vaccine should elicit a large response to this antigen. This requires that the antigen rapidly elicits immunity (i.e. has low *φ_X_*, and in terms of immunology it should be an immunogenic antigen), and also requires a high vaccine viral load to generate a large response. Engineering this requires tackling a trade-off between avoiding vaccine clearance (i.e. having a low *k_X_*) but allowing for rapid clearance of the pathogen (having a high *k_P_*). Vaccines designed to express the antigen in a form that is different from that in the pathogen might help solve this problem. Thus, to elicit immunity to influenza, one might design secreted forms of the hemagglutinin or neuraminidase proteins. A recombinant hemagglutinin protein that is secreted rather than on the virion surface would prevent the antibody response to this protein from clearing the recombinant vector vaccine (have low *k_X_*) without compromising the clearance of the influenza virus pathogen which has hemagglutinin on its surface (i.e. has high *k_P_*). In this manner our model allows the identification and tuning of parameters that affect vaccine efficacy, and a comprehensive search of parameter space would identify ideal combinations of vaccine properties. We now turn to vaccine designs that overcome problems created by evolution, our specific interest here.

#### 4.3.1 Control the inoculum

The results above suggest that vaccine evolution is only likely to compromise immunity to the antigen if there is substantial evolution and this evolution results in more rapid clearance of the vaccine virus. In this case, one possible solution takes advantage of the short-term nature of vaccine growth: control the inoculum. Two ways of controlling the inoculum are to control its composition and to control its size. Evolution can be reduced by purifying the inoculum - an inoculum that is entirely vaccine cannot begin to give way to revertant until some are generated by mutation, hence a low (or zero) density of revertant in the inoculum enhances the duration of within-host vaccine utility. If it not feasible to *eliminate* the revertant from the inoculum, it can nevertheless be beneficial to lower the frequency of the revertant virus in the inoculum. The effect of revertant frequency in the inoculum is evident in Figure 6: the magnitude of immunity to the vaccine increases by orders of magnitude as the initial frequency of the revertant is decreased.

**Figure 6:**
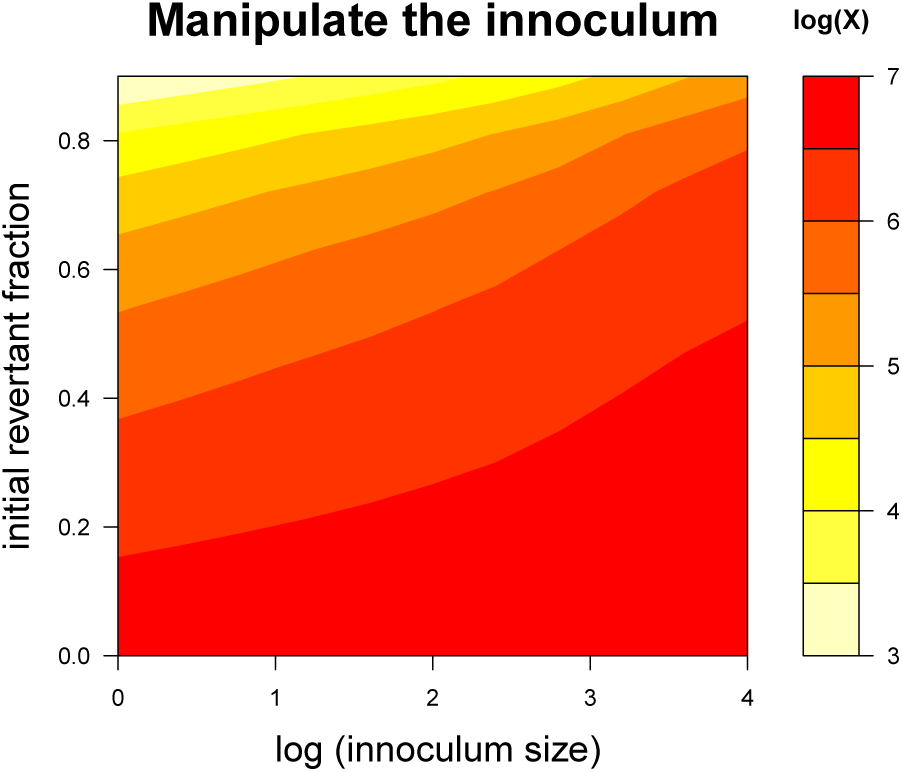
Effects of manipulating the inoculum on immunity to the vaccine. Small inocula that contain vaccine plus revertant are more prone to reduced immunity levels than are large inocula with little revertant. Composition of the vaccine has the larger effect for these parameters, as indicated by the contours being more horizontal than vertical. An intrinsic fitness cost of *c* = 0.1 was set for these trials. Smaller *c* values would lead to higher vaccine and immunity levels across the graphs.

Evolution can also be reduced by increasing the inoculum size. To achieve a threshold antigen level, a large inoculum requires less growth than a small one. Less growth means less evolution – in the extreme, a large enough inoculum requires no vaccine growth, as with killed vaccines. Figure 6 also shows the consequences of changes in inoculum size. When the revertant frequency in the inoculum is high, increasing the inoculum size appreciably increases the magnitude of immunity; a much reduced benefit is seen when revertant frequency is low, likely because there is less evolutionary interference from the revertant. These results hint at a potential tradeoff between the benefits of reducing the frequency of the revertant in the inoculum and increasing the dose. Consideration of this tradeoff could help choose an economically feasible strategy, since both purifying the inoculum and increasing its dose are likely to incur financial costs.

Whether and how well controlling the inoculum will work in practice will depend on details. Solutions may be quantitative rather than absolute. Intuition is useful for guidance but needs to be confirmed by formal analyses, guided by data for the specific implementation.

#### 4.3.2 Design the vaccine

Controlling the inoculum corrects the evolution problem by circumventing the consequences of vaccine inferiority. A different solution is to design the vaccine with less of a disadvantage. The most obvious realm for this approach is in vaccine engineering: the timing and tissues of antigen expression, location of the transgene in the vector genome, and the size of the transgene may all influence intrinsic fitness effects [9,10,17,38,39]. Directed evolution approaches might also work: one simple approach in reducing an intrinsic cost might be to ‘pre-adapt’ the vector *in vitro* on host cells expressing the antigen *in trans*. This adapted vector would then be used as the vaccine backbone. Another simple approach would be to compete several different vaccine designs *in vitro* and pick the design with fastest growth. Any approach using *in vitro* adaptation needs to avoid adapting the vector to the extent that it compromises ability to grow *in vivo*.

## 5. Discussion

Any live viral vaccine may evolve within the host. The potential for attenuated viruses to revert to wild-type virulence is well appreciated [1,2], even if it presents a problem for relatively few vaccines (e.g., attenuated polio: [40]). There is also a potential for live, recombinant vector vaccines to evolve – our focus in this paper – with the main concern being loss or reduced expression of the transgenic insert [4,41]. If the vaccine evolution occurs fast enough or the vaccine infection persists long enough, loss of the insert could reduce vaccine efficacy.

We developed and analyzed models to explore ways in which vaccine evolution could lead to a reduction in vaccine efficacy. An intrinsic fitness advantage of the revertant virus, expected because engineering transgene expression is likely to have metabolic and other costs, will lead to vaccine being gradually overgrown by revertant. Yet this is only likely to cause a reduction in the immunity to the vaccine antigen if it leads to a reduction in the absolute amount (as opposed to merely a reduction in relative frequency) of the vaccine virus. Ascent of revertant can reduce the amount of the vaccine virus if the revertant uses resources required for virus replication or the vaccine virus is cleared by the innate or adaptive responses elicited by the revertant.

Our results revealed that that for a broad parameter regime, within-host evolution is unlikely to cause a significant loss of vaccine efficacy (i.e. reduction in the level of immunity to the inserted transgene). Furthermore, undesirable consequences of vaccine evolution may often be easily remedied by ensuring the frequency of the revertant virus in the inoculum is low and by increasing the size of the inoculum. We also suggest that further gains in vaccine efficacy can be achieved by appropriate engineering of the vaccine antigen, allowing it to elicit immunity that clears the pathogen but not the virus vaccine, although such engineering may not be easy.

One major outcome of our analysis was that intuition about vaccine evolution was not easily translated into intuition about immunity. Indeed, even intuition about evolution often failed because that intuition was based on vaccine versus revertant fitness, but the vaccine growth phase was short enough that differential fitness had little effect on evolution. Even more fundamentally, intuition sometimes failed because the development of immunity to vaccine could be unaffected by the revertant. Thus, our intuition suggested that vaccine inferiority could stem from both an intrinsic fitness disadvantage and a disadvantage due to adaptive immunity to the transgene/antigen. Both effects were found to impair the development of immunity to vaccine, but not necessarily for the reasons suggested by our intuition.

Measuring the intrinsic fitness effect of the transgene is likely to be an important step in vaccine design. For assessing vaccine evolution, the relevant biological realm is within the host. Nonetheless, *in vitro* growth environments may reveal much about a vaccine’s intrinsic propensity to evolve loss of antigen expression. There are various ways intrinsic fitness effects and their evolutionary consequences might be studied. Vaccine growth in tissue culture may reveal some aspects of intrinsic fitness effects and should be relatively easy to study. Loss of the transgene *per se* would be detectable by PCR, and the fitness advantage of revertant over vaccine could be measured from changes in revertant frequency. The quantitative relevance of an *in vitro* estimate to *in vivo* growth would be unknown, but the measure should allow qualitatively comparing engineering designs that improve intrinsic vaccine fitness. If vaccine reversion were due to down regulation of the transgene instead of loss, fitness estimation would require knowing the mutations responsible and monitoring their frequencies. Use of culture-wide antigen levels to measure fitness might provide a sense of whether vaccine evolution would lead to reduced antigen levels *in vivo*, but it would be less sensitive in measuring evolution than is measuring mutation frequencies.

*In vitro* assays may be useful in measuring intrinsic fitness effects, but *in vivo* – in the patient – is the ultimate environment for studying within-host evolution and its effects. Not only are the dynamics of viral spread different between *in vitro* and *in vivo* environments, but most immune components will be in play only *in vivo*. Furthermore, those components may vary across tissues within the host. Sampling across this heterogeneity *in vivo* will be challenging but may be necessary to know whether, when, and where vaccine evolution is a problem. If revertant remains a minority of the population, we expect that vaccine evolution can be ignored. Perhaps *in vitro* studies of vaccine evolution will provide most of the information relevant to *in vivo* evolution, but it is too early to know.

We have focused on recombinant vector vaccines that cause acute infections. Necessarily, our recommendations are based on simple models that are caricatures of the complex within-host dynamics of acute infections. Simple models are appropriate at this stage because of uncertainties at many biological levels, and under these circumstances simple models frequently generate more robust results than do complex models [42,43].

The generation of innate and adaptive responses can be modeled with different assumptions than used here, and those alternative processes may affect the conclusions. For example, time-lags in the activation of cells may dominate the time for the generation of an innate immune response, with virus density having a consequently smaller role than assumed here (as can be seen in [44] and modeled in [29]). We have modeled that responses to different antigens are generated independently of each other and do not compete. We have done so because vaccines are likely to cause relatively mild infections during which the densities of pathogen and immune cells do not reach sufficiently high levels required for competitive interactions to be important. The adaptive immune response may be more influenced by recruitment which is followed by a period of proliferation even in the absence of antigen [45–47]. Both these scenarios would minimize the impact of evolutionary changes in the vaccine on the amount of immunity generated to the transgene.

Finally, it is easily appreciated that there are realms we do not consider, such as spatial structure [48] and recombinant vector vaccines based on viruses such as Cytomegalovirus that cause persistent infections [49] or that are transmissible. Spatial structure may limit the impact of vaccine evolution on immunity (e.g., prevent mutants from taking over the entire population). In contrast, vaccines that cause persistent infections or are transmissible are likely to be more severely affected by evolution than are vaccines causing acute infections, as there is a longer timeframe for evolution to operate.

With so little experience from recombinant vector vaccines, we can merely guess how commonly within-host evolution will compromise vaccine efficacy. Given that simple steps can be taken to reduce vaccine evolution, vaccine development programs should at least entertain the possibility that evolution can underlie failure. Avoiding vaccine evolution may be easier than developing an entirely new vaccine.

## Appendix: the models

The models used here specify features of viral infections. Some basics include the following:

1. Two viral types: Only vaccine and wild-type (vector or revertant) are ever present.
2. Acute infections. Infections are short term because they are subject to control and clearance by any combination of three factors: resource limitation, innate immunity and adaptive immunity. Further details not included in this Appendix can be found in Supplements.

### 1. Model formulation

#### Variables

The following table defines the variables used in these equations.

| Term | Description |
| --- | --- |
| $V$ | Vaccine |
| $W$ | Wild-type (revertant) vector |
| $R$ | Resource |
| $X$ | Adaptive immune response specific to recombinant (vaccine) antigen |
| $Y$ | Adaptive immune response to vector/revertant as well as to vaccine |
| $Z$ | Innate immune response (common to vaccine and revertant) |

#### Parameters

The following table gives the parameter definitions and values used, except where different values are specified in figures or supplementary material.

| Abbreviation | Description | Values |
| --- | --- | --- |
| Intrinsic factors |  |  |
| $r$ | rate of growth of $V$ | 3 per day |
| $c$ | cost to having recombinant antigen | $0 < c < 1$. |
| $\mu$ | mutation rate for $V \rightarrow W$ | $0, 10^{-3}, 10^{-6}$ |
| $d$ | death rate of virus | 1 |
| Resource limitation |  |  |
| $\phi_R$ | resource for half max growth of $W$ and $V$ | 1 |
| Innate immunity |  |  |
| $\sigma$ | rate of stimulation of innate immunity | $2.7 \times 10^{-5}$ |
| $k_Z$ | killling rate of V due to Z (innate immunity) | $3 \times 10^{-2}$ |
| $d_Z$ | decay of innate immunity in absence of antigen | 1 [0 for model 2] |
| Adaptive immunity |  |  |
| s | rate of clonal expansion of adaptive immunity | 3 |
| $\phi_X$ | antigen for half max growth of adaptive immunity X | $10^3$ |
| $\phi_Y$ | antigen for half max growth of adaptive immunity Y | $10^3$ |
| $k_X$ | killling rate of V due to X (immunity to insert) | $10^{-6}$, various |
| $k_Y$ | killling rate of V & W due to Y (immunity to vector) | $10^{-6}$, various |
| Initial conditions |  |  |
| $X(0)$ | initial immunity to vaccine antigen | 1 |
| $Y(0)$ | initial immunity to vector antigen | 1 |
| $Z(0)$ | initial innate immunity | 0 |
| $V(0)$ | initial virus | 1 |
| $W(0)$ | initial revertant | various |
| $R(0)$ | initial resource | $2 \times 10^5$, $10^{20*}$ |

\*The higher value of $R(0)$ is used to eliminate it as a basis for viral clearance.

#### Equations

Resources start with a fixed amount and are depleted by vaccine and revertant growth, without replenishment:

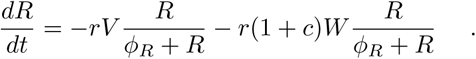

The vaccine virus grows on resource R at rate r, depleted by mutation, death, and all 3 types of immunity:

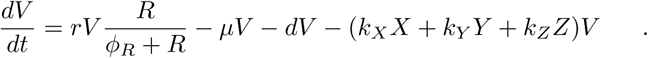

Revertant grows on resource R at rate r(1+c), depleted by mutation, death, and 2 types of immunity (not *X*):

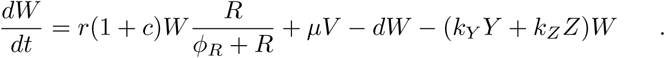

Adaptive immunity specific to vaccine grows according to its present value and a discounted value of the current vaccine density:

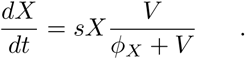

Adaptive immunity common to vaccine and revertant grows according to its present value and a discounted value of the current vaccine plus revertant densities:

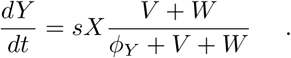

Innate immunity, also common to vaccine and revertant, grows according to current levels of vaccine and revertant, with diminishing growth as a limit is approached. Innate immunity also decays:

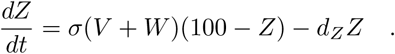

These models follow the usual assumptions of SIR models, except that susceptible hosts (host cells in our case) are modeled as Resource. As is typical in these models, variables for ‘free’ virus are omitted, an assumption based on the quasi-steady state approximation (Perelson 2002).

